# Remote activation of place codes by gaze in a highly visual animal

**DOI:** 10.1101/2024.09.30.615982

**Authors:** Hannah L. Payne, Dmitriy Aronov

**Affiliations:** Zuckerman Mind Brain Behavior Institute, Columbia University; Howard Hughes Medical Institute at Columbia University

## Abstract

Vision enables many animals to perform spatial reasoning from remote locations. By viewing distant landmarks, animals recall spatial memories and plan future trajectories. Although these spatial functions depend on hippocampal place cells, the relationship between place cells and active visual behavior is unknown. Here, we studied a highly visual animal, the chickadee, in a behavior that required alternating between remote visual search and spatial navigation. We leveraged the unique head-directed nature of avian vision to track gaze in freely moving animals. We discovered a profound link between place coding and gaze. Place cells activated not only when the chickadee was in a specific location, but also when it simply gazed at that location from a distance. Gaze coding was precisely timed by fast, ballistic head movements called “head saccades”. On each saccadic cycle, the hippocampus switched between encoding a prediction of what the bird was about to see and a reaction to what it actually saw. The temporal structure of these responses was coordinated by subclasses of interneurons that fired at different phases of the saccade. We suggest that place and gaze coding are components of a unified process by which the hippocampus represents the location that an animal is currently attending to. This process allows the hippocampus to implement both local and remote spatial functions.

## MAIN

Consider a classic example of spatial memory: an animal remembering a nut hidden at the base of a tree. Theories of hippocampal function posit that such a memory depends on place cells – in this case, the set of neurons active at the base of the tree^1,2^. Therein lies a problem: to efficiently find the tree and retrieve the nut, the animal must first recall the memory from a remote location, where a completely different set of place cells might be active. How are place cells compatible with such a remote function of the hippocampus? In visual animals, remote recall is often driven by gaze^3^. By simply looking at a place from a distance, animals can recall associated information without physically revisiting that location. Yet, it is unknown how the brain coordinates remote vision with the activity of place cells.

This problem is unsolved largely because eye tracking in freely behaving animals is extremely challenging. In addition to this technical issue, many laboratory models, including rodents, lack foveal vision^4^ and rarely orient their eyes precisely at visual targets. It is often impossible to know exactly what these animals are looking at, even when eye tracking is feasible^5–8^. In primates that have foveal vision, hippocampal activity does represent gaze targets^9–14^ and other visual information^15–17^. However, these animals are usually recorded stationary, or in conditions where it is hard to disambiguate gaze coding from place coding.

To address these challenges, we leveraged a unique feature of avian biology. Birds, like primates, have foveal vision and actively control their gaze to fixate visual targets^4,18,19^. However, instead of eye movements many bird species rely primarily on head movements to shift gaze from one target to another. These head movements are much more feasible to track in a small, freely moving animal. We chose to use the black-capped chickadee – a member of a food-caching bird family that has abundant place cells in the hippocampus^20,21^. Chickadees provide a unique opportunity to simultaneously study spatial coding and gaze during unconstrained behavior.

### Head-directed gaze strategies in chickadees

All birds use their heads to direct gaze, but the nature of head movements and the extent of additional eye movements are highly variable across species^4,22^. We therefore started by characterizing these behaviors in chickadees. For head tracking, we adapted a multi-camera system^23^ that triangulated infrared-reflective markers on the bird’s head (Fig. 1a). As in other species^4,19^, these measurements revealed a saccadic-like behavior. Chickadees produced fast, ballistic changes in head angle (“head saccades”) interleaved with periods of stable head angle (“head fixations”). Head saccades were 76 ± 21 ms in duration, occurred at instantaneous rates of 3.8 ± 1.4 Hz, and were in several additional ways remarkably similar to eye saccades in primates^3^ (mean ± standard deviation, n = 3.3×10^5^ saccades in 8 birds; Fig. 1c, Fig. S1a-c).

**Figure 1.**
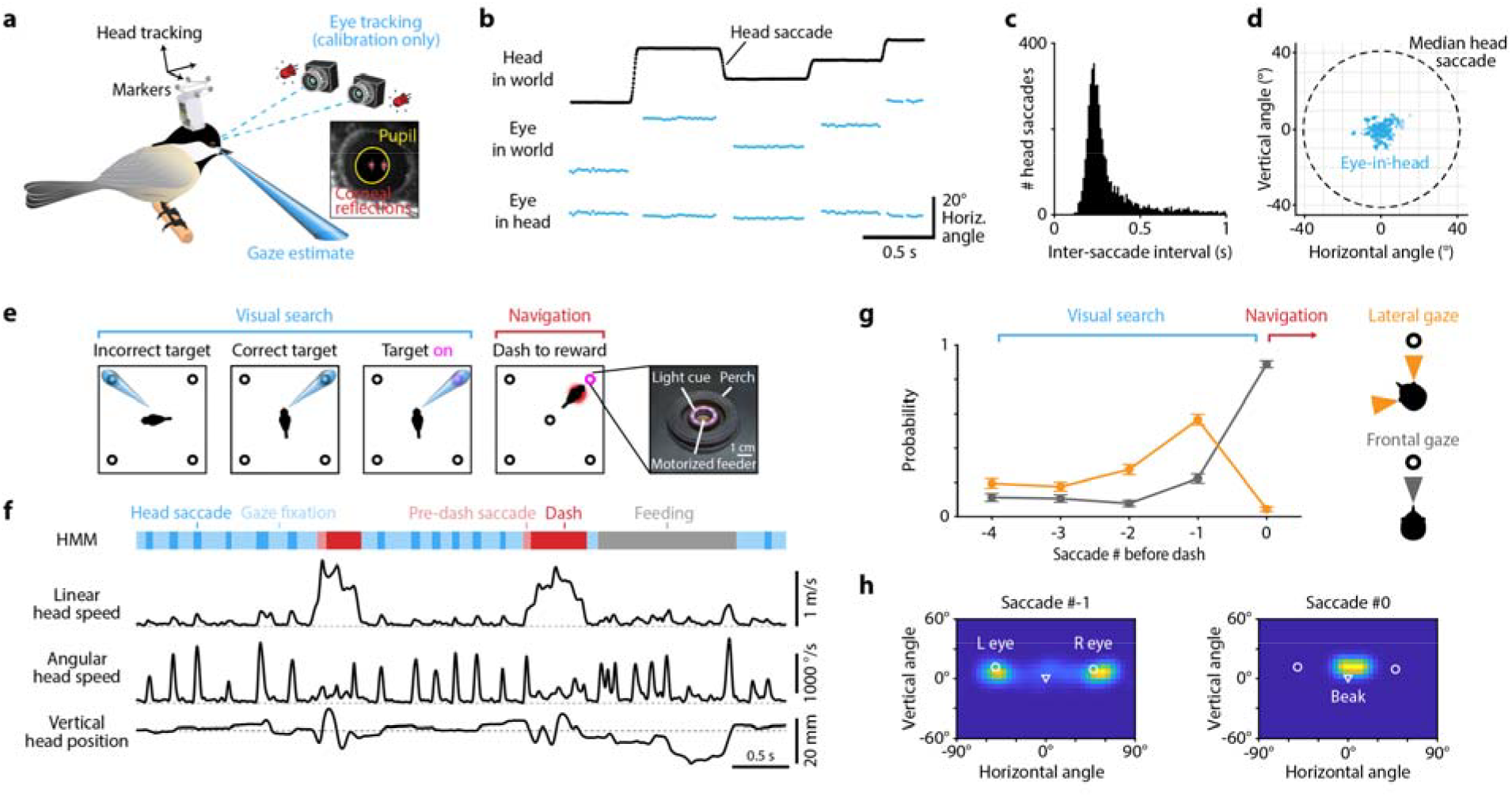
Head-directed gaze strategies in chickadees. **(a)** Head position is tracked using four infrared cameras (not shown) and reflective markers on the head. In a calibration experiment, eye position is measured using two additional cameras that track the pupil and corneal reflections of two light sources. **(b)** Head angle shows prolonged fixations interrupted by fast, ballistic movements (“head saccades”). For simplicity, only the horizontal angle is plotted. The eye could not be accurately tracked during head saccades; those points are omitted. When head position is subtracted from eye position, the residual eye position relative to the head shows very little movement. **(c)** Distribution of intervals between head saccades from a single session. **(d)** Distribution of eye positions from an example calibration experiment. Eye movements relative to the head are much smaller than the head saccades. **(e)** Schematic of the discrete visual search task, which contains a visual search period and a separate navigational period. Gaze at the correct target activates a light cue. The chickadee then navigates to that target to obtain a reward. **(f)** Different behaviors involve distinct movements of the head and can be segmented from head tracking data using a Hidden Markov Model (HMM). **(g)** Time course of the bird’s use of two different gaze strategies (lateral and frontal gaze). Saccade #0 is defined as the one immediately preceding a dash (“pre-dash saccade” in (f)). Data are mean ± standard error of the binomial proportion for one example session (n = 250 dashes). **(h)** Distribution of head orientations relative to the target, averaged across all #0 and #-1 saccades for the same bird as in (g). Symbols mark the orientation of the two pupil centers and the tip of the beak. Saccade #0 was usually frontal (using both eyes). Prior to that, saccades tended to be lateral (using one eye at a time).

We then performed a calibration experiment in which we tracked eye movements in addition to head movements. For this purpose, we engineered a dual-camera video-oculography system that estimated the pupillary axis using corneal reflections of two infrared light sources (Fig. 1a, Fig. S1d-f). These measurements were possible only when we encouraged chickadees to perch directly in front of the cameras. We found very little movement of the eyes relative to the head: 5.4 ± 0.4° median absolute deviation, nearly an order of magnitude smaller than the movement of the head itself (n = 8 birds, Fig. 1d, Fig. S1g). We concluded that chickadees almost exclusively rely on head saccades to direct gaze during free motion. Head tracking is therefore sufficient to determine where a bird is looking, provided that gaze targets in a behavioral task are separated by more than ∼10°. Unlike eye tracking, which requires the head to be nearly stationary relative to the camera, head tracking is feasible in chickadees freely navigating across a behavioral arena.

To study place and gaze coding, we trained birds on a discrete visual search task (Fig. 1e). Each trial began with the chickadee perched in the center of the arena, choosing between four equidistant sites that consisted of a perch, a light cue, and a motorized feeder. We first ran a simple version of the task to characterize the chickadees’ visual search behaviors. In this version, there was a delay period of random duration, followed by a light cue that indicated one randomly chosen site. The bird could approach the indicated site to retrieve a piece of a sunflower seed. Chickadees typically approached rewarded sites via fast, direct movements that we call “dashes”. We used a Hidden Markov Model to segment dashes, head fixations, saccades, and feeding periods in the head tracking data (Fig. 1f).

We found that chickadees used two distinct gaze strategies in our task (Fig. 1g-h, Fig. S1h). During the delay period, they shifted gaze between different sites using one eye at a time (“lateral gaze”). In other words, chickadees oriented their head to align either the left or the right pupil with one of the sites. Because the light cue was intentionally dim, chickadees probably used this behavior to search for the rewarded site using the foveal region of the retina^18^. After locating the light, birds instead oriented the beak toward the target (“frontal gaze”), viewing the rewarded site with the non-foveal region of both eyes. They usually followed this frontal gaze by a dash. Other bird species have similar strategies, using lateral gaze to investigate objects and frontal gaze during directed movements^24–26^.

With these results in mind, we modified the task to create a closed-loop version. Instead of enforcing a delay period, we activated the light cue in response to the bird gazing at the correct site. Because chickadees had no *a priori* knowledge of which site was rewarded, they often gazed at multiple incorrect sites before choosing the correct one. The closed-loop structure allowed us to precisely control the timing of the visual stimulus relative to the saccade. It also ensured that the cue was not detectable by peripheral vision during off-target saccades; this feature will become important later. In this closed-loop task, we performed recordings of the anterior hippocampus using silicon probes.

#### Place cells are activated by remote gaze

We measured place tuning and gaze tuning in the firing of hippocampal neurons. To match published work^27,20,21^, we analyzed place tuning only during periods when the chickadee was locomoting – i.e., dashing between sites (Fig. 2a). In contrast, we analyzed gaze tuning during stationary visual search periods, when the bird was saccading and fixating at targets (Fig. 2b). We started by examining activity as a function of contralateral gaze – i.e., the gaze target of the eye contralateral to our hippocampal recording. As in other behavioral tasks, many hippocampal neurons in chickadees were place-tuned: of the 1929 putative excitatory cells in seven birds, 62% were classified as place cells. We found that a similarly large fraction of cells were gaze-tuned: 57% of the same 1929 neurons. Many of these cells had strong gaze fields that were tightly localized in the environment, qualitatively similar to conventional place fields (Fig. 2c-d).

**Figure 2.**
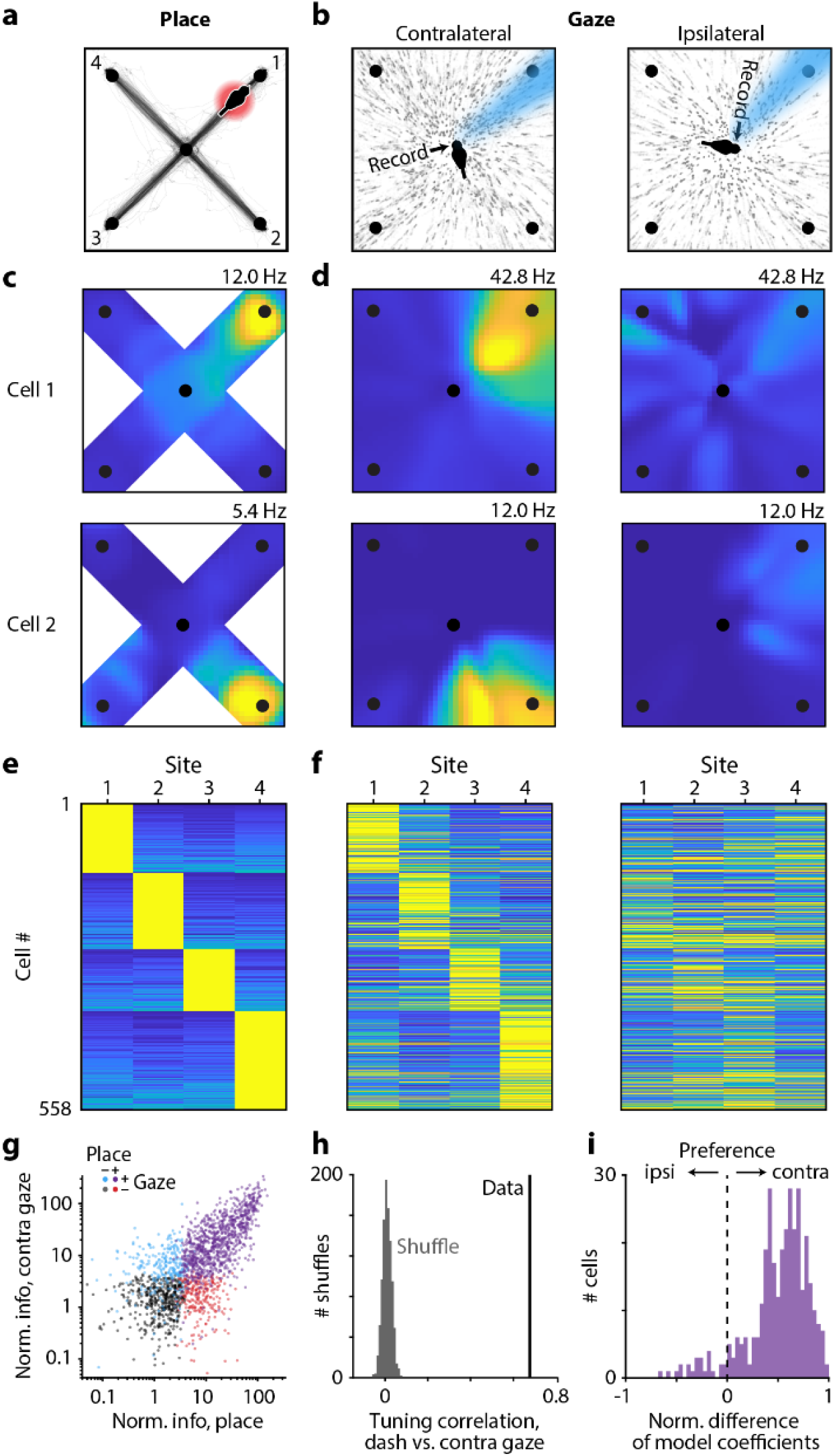
Place cells are activated by remote gaze. **(a)** Place representation is measured during times when the bird is dashing toward a target. Black trace: trajectory of the bird during such periods in an example session. **(b)** Gaze representation is measured during times when the bird is saccading from the center of the arena. Black dots: projected gaze in the example session, plotted separately for the eyes contralateral and ipsilateral to the hippocampal recording. **(c)** Mean firing rate as a function of the bird’s location for two example place cells. Only places on the X-shaped part of the arena are shown because the chickadee almost never visits other locations. Color scale is from 0 (blue) to the indicated maximum (yellow). **(d)** Mean firing rate as a function of gaze location for the same two cells. **(e)** Peak firing rate during dashes to each of the four target sites. Excitatory place cells with strong selectivity for one of the targets (>0.5 selectivity index) are shown. Cells are sorted first by location of their strongest response and then by the magnitude of their second-strongest response. Each row is normalized to the maximum. **(f)** Coefficients of a model that fits gaze responses as a combination of tuning to contralateral and ipsilateral gaze. Cells and sorting are the same as in (e). Rows are normalized to the maximum across both matrices in (f). **(g)** Strength of place and gaze tuning across all excitatory cells, colored based on a statistical threshold for place and/or gaze tuning. “Normalized information” is mutual information between spikes and the behavioral variable (place or contralateral gaze, discretized into four target sites), divided by the mean for shuffled data. The two types of tuning are strongly correlated. **(h)** Correlation of tuning curves for place and contralateral gaze (r = 0.68, n = 558 cells). Tuning curves are measured across the four targets for all place cells shown in (e,f). Shuffled distribution is obtained by scrambling cell identities. **(i)** Comparison of contralateral and ipsilateral gaze responses, using model coefficients in (f) for each cell’s preferred dash target. Included cells are those in (h) with >0.5 selectivity both for place and for gaze with at least one of the eyes (n = 289 cells). Normalized difference is (c-i)/(c+i), where c and i are contralateral and ipsilateral coefficients. Gaze responses are almost entirely explained by contralateral tuning.

There are other known situations in which the hippocampus encodes more than one experimental variable^2,17,20,28,29^. In these cases, different kinds of selectivity are usually mixed randomly within the recorded population. To our surprise, this was not true for place and gaze tuning. The amounts of information encoded about place and about gaze were strongly correlated across cells (Fig. 2e-g). In fact, 75% of all place cells were also significantly tuned to gaze, compared to 57% expected from random mixing.

Not only did the same neurons encode place and gaze, but these two representations had a striking overlap in space (Fig. 2c,d). To quantify this overlap, we selected place cells that had a strong preference for dashes toward a single site. Tuning curves for place and gaze were strongly correlated in these cells (Fig. 2e,f,h). In those cells that also had a strong preference for a single gaze target, the preferred place was the same as the preferred gaze in 95% of cases, compared to ∼25% expected by chance. These analyses show that place and gaze tuning are not represented independently in the hippocampus. Rather, remote gaze activates place representations. In other words, a place cell is active not only when a bird is physically in a certain location, but also when the bird simply looks at that location from a distance.

In many behaviors, the hippocampus can represent the future location of the animal^30,31^. In our task, birds often look at sites before visiting them. Therefore, a conceivable explanation of our results is that the hippocampus actually represents future location, which happens to correlate with gaze. To determine whether gaze responses were specifically related to visual behavior, we relied on a unique feature of avian anatomy. As in mammals, the avian hippocampus sits at the top of the visual hierarchy and receives input from multiple visual pathways^32–34^. However, in most birds, the optic tract fully crosses to the contralateral hemisphere at the optic chiasm^35^. There is also very limited cross-hemispheric communication in the visual system due to the lack of a corpus callosum. As a result, visual functions are highly lateralized^36,37^. If gaze signals are actually driven by future location, we should observe them bilaterally, but if they are specific to gaze, we might expect them to be lateralized in the hippocampus.

We found place and gaze tuning in both hemispheres. However, gaze tuning was evident only when we analyzed the eye contralateral to the recorded hippocampus (Fig. 2d). Neurons responded only when the contralateral, but not the ipsilateral, eye looked at the preferred target. To illustrate this result across the population, we implemented a model that fit neural activity as a combination of tuning to ipsilateral and contralateral gaze. Such a model was necessary because chickadees sometimes gazed simultaneously at two sites with different eyes. In this model, activity was almost entirely explained by contralateral gaze (Fig. 2f,i). This result was true regardless of whether birds were allowed to use either eye or only the contralateral eye to trigger the reward (Fig. S2). We concluded that gaze tuning is truly specific to where the bird is looking.

Finally, one consideration is that chickadees in our task always performed the visual search from the same central location of the arena. Our recordings are therefore consistent with hippocampal activity encoding the direction of gaze, rather than the location of the gaze target. To test this possibility, we trained three birds on an “all-to-all” task in which they dashed directly between five target sites without returning to a central perch (Fig. S3). We found that gaze responses were more invariant to the site from which the chickadee was gazing (the “source”), than to the site at which it was looking (the “target”). There were some differences in firing between gaze sources to the same target. But the same differences were also observed between dashes from different sources, consistent with known examples of trajectory remapping in the hippocampus^38,39^. We concluded that hippocampal responses encoded the location of the gaze target, rather than simply the direction of gaze.

#### Gaze responses encode an internal prediction

An overarching question of our study is whether the hippocampus can recall internal information about the visual world, such as memories associated with specific viewed locations. For this purpose, it is not sufficient to simply react to visual stimuli; rather, neural responses should contain additional, internally-driven information about gaze targets. To test this idea, we first asked whether the timing of hippocampal activity was consistent with purely sensory responses. We aligned neural activity to the time of peak angular head velocity during each saccade. We compared these responses to the timing of activity during dashes. Surprisingly, we found that saccade-aligned activity, but not dash-aligned activity, was biphasic (Fig. 3a-d). Neurons produced the first peak in firing during the saccade itself at 17 ± 14 ms, and then the second peak at 187 ms ± 4 ms, roughly corresponding to the time of the next saccade (n = 278 cells, mean ± bootstrapped standard error; Fig. 1c). Most neurons participated in both phases of the response, though the relative amplitudes of the two peaks varied across cells.

**Figure 3.**
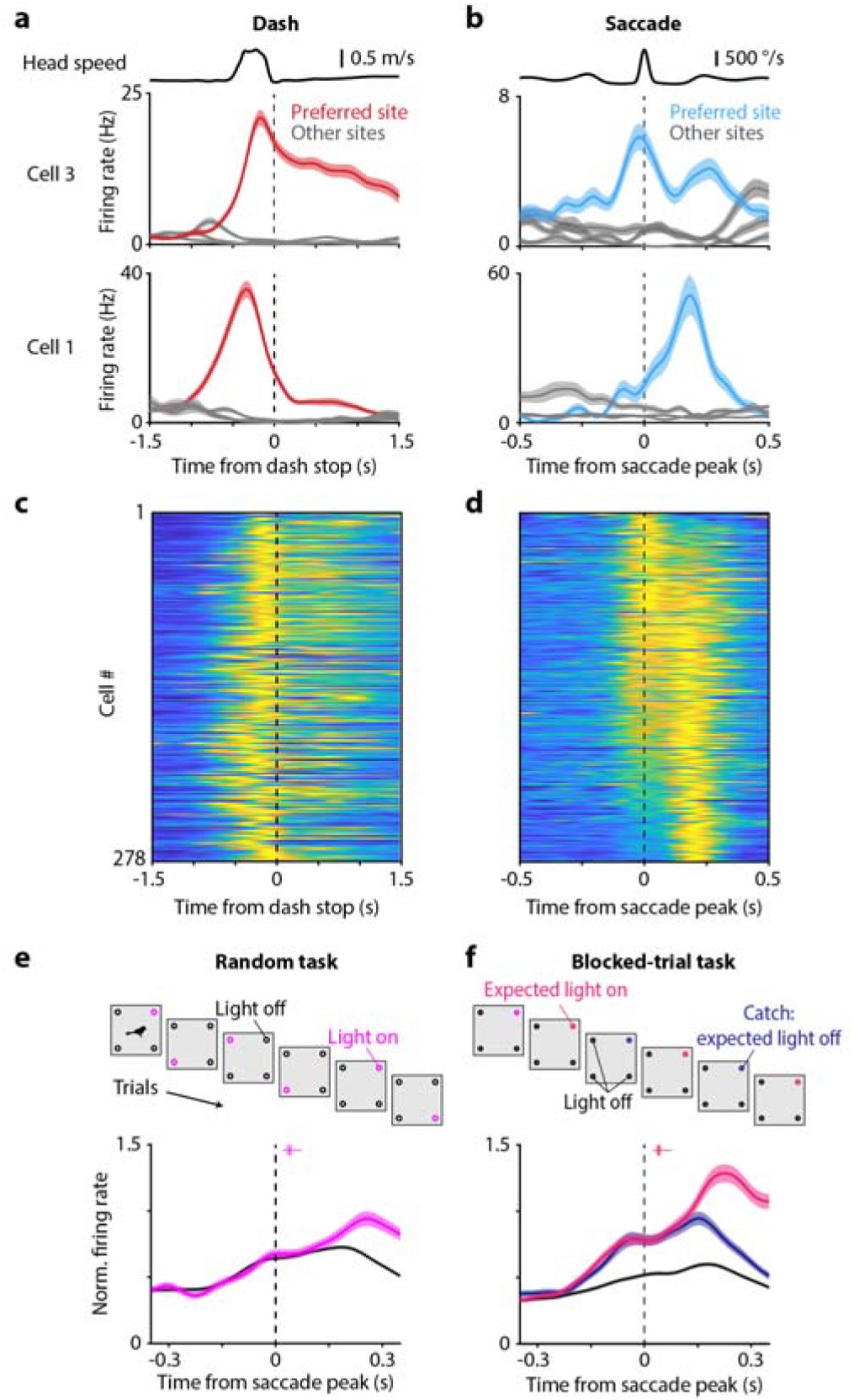
Gaze responses encode an internal prediction. **(a)** Activity of two excitatory cells aligned to dashes toward each of the four targets. Average linear speed of the head across the entire dataset is shown above. **(b)** Activity of the same two cells aligned to saccades toward the same four targets. Average angular speed of the head is shown above. **(c)** Activity of cells aligned to dashes toward their preferred target. Included are excitatory cells that have strong selectivity (>0.5) of both place and gaze responses for the same target. Each cell’s activity is normalized from 0 (blue) to its maximum (yellow). **(d)** Activity of the same cells aligned to saccades toward their preferred target. Cells are sorted by the difference in firing during the early and the late phases of the response (centered on 17 and 187 ms); the same sorting is applied to (c). **(e)** Responses to the preferred gaze target in the Random task, separately for saccades when that target was correct (indicated by the light turning on) vs. incorrect (indicated by the light staying off). Included are excitatory neurons with significant place and contralateral gaze tuning, peak firing rate of >1 Hz in the Light-off condition, and at least 5 saccades to the preferred gaze target (n = 271 cells); each cell’s activity is normalized to the peak rate during saccades in the Light-off condition. Box and whisker: 5^th^, 25^th^, 50^th^, 75^th^, and 95^th^ percentiles of the latency to the light turning on (median = 40 ms). **(f)** Responses to the preferred gaze target in the Blocked-trial task, separately for saccades when the chickadee expected the target to turn on vs. stay off (n = 402 cells; median latency to light on 40 ms). The former category includes catch trials, when the target was expected to turn on, but stayed off.

In other words, a neuron tuned to a specific target location was active twice: once at the start of a visual fixation on that target (“early response”), and once at the end of the fixation (“late response”). The late response was compatible with latencies expected in the visual system. In contrast, the early response was certainly not visual: it occurred largely before the chickadee fixated on the target and started even before the head began to move. Yet, this early response had the same gaze tuning as the late response (Fig. 3b). In other words, early activity was tuned not to the current gaze location, but to the gaze location of the future fixation.

These results suggest an intriguing hypothesis: the early response might encode an internal prediction of what the bird is about to see, while the late response might encode what the bird just saw. Thanks to the closed-loop design of our task, we could try to separately influence these two responses. We analyzed each cell’s responses to its preferred gaze target. We compared saccades when that target site was rewarded (and the light cue turned on) to saccades when the same site was unrewarded (and the light cue stayed off; Fig. 3e). In the early response, firing rates were identical between these two conditions; this was expected because the hippocampus had no *a priori* information about the upcoming light cue. In contrast, firing rates diverged in the late response, with higher rates in the light-on condition. We concluded that late in the fixation period, the hippocampus responds to visual stimuli. This late response can represent information beyond the location of the gaze target.

Next, we asked how hippocampal responses changed if the chickadee was able to predict the upcoming light cue. Instead of rewarding a random site on each trial, we implemented a blocked-trial task in three birds, in which the same site was rewarded six trials in a row. Chickadees found the rewarded site faster during these trial blocks, suggesting that they understood the structure of the task (Fig. S4). We found that in the blocked-trial task, firing rates diverged during the early response (Fig. 3f). In other words, the early response was predictive of the light cue, even before the chickadee gazed at the correct site.

After diverging in the early response, firing rates continued to separate during the late response. Here, we wanted to disambiguate the bird’s prediction from the actual reaction to the light cue. We included a small number of “catch trials” in which the reward was omitted – i.e., the chickadee expected the light to turn on, but the light actually stayed off. Absence of the expected light cue suppressed neural activity: the late response was weaker in catch trials compared to light-on trials. However, this response was still stronger than in those trials when the chickadee did not expect the light cue.

In summary, the early response during saccades seemed to represent the chickadee’s prediction of an upcoming visual stimulus. In contrast, the late response represented a mixture of the prediction and the reaction to the actual stimulus. Note that the “early” response relative to one saccade coincides in time with the “late” response relative to the previous saccade. Similarly, the late response coincides with the early response of the next saccade. Therefore, another way to describe our data is that activity during each saccade represents a mixture of information about the recently completed visual fixation and the upcoming fixation. We are unsure of what exactly the hippocampus represented in our task: it could be simply the visual stimulus, or something more complex like reward anticipation. Regardless, the important conclusion is that the hippocampus represents both internally-driven and externally-driven information about visual targets. This coding is temporally coordinated by saccadic head movements, multiple times per second.

#### Inhibitory dynamics during saccades

Saccade-related dynamics that we observed are not unlike other fast phasic processes in the hippocampus – most notably the theta oscillation. Inhibition plays a major role in these processes. For example, different classes of inhibitory interneurons fire at different phases of theta^40^. Such precise timing is thought to be important for the temporal patterning of excitatory cells and for the mechanisms of synaptic plasticity^41,42^. Birds do not appear to have continuous oscillations in the hippocampal local field potential (LFP)^20^. We wondered whether their inhibitory (I) and excitatory (E) cells were instead temporally coordinated by head saccades.

As in previous studies^42,20,21^, we classified chickadee hippocampal neurons as E or I using firing rates and spike waveforms (Fig. S5). Up until this point, we have been reporting only the activity of E cells, but now consider I cells. Our analysis revealed two types of gaze responses in I cells. Some neurons (e.g. cell 1 in Fig. 4a) produced a smaller trough in firing shortly after the saccade and a larger peak in firing later during the fixation. Other neurons (e.g. cell 2 in Fig. 4a) instead produced a smaller peak early and a larger trough later. We summarized these patterns by measuring the instantaneous phase of each cell’s response at a fixed time after the saccade. Across the population, these phases had a clearly bimodal distribution, with two groups of cells roughly 180° apart (Fig. 4b-d). These groups (“Peak” and “Trough” cells) also had different mean firing rates and spike waveforms (Fig. 4d, Fig. S5). We concluded that Peak and Trough cells likely correspond to different classes of hippocampal interneurons.

**Figure 4.**
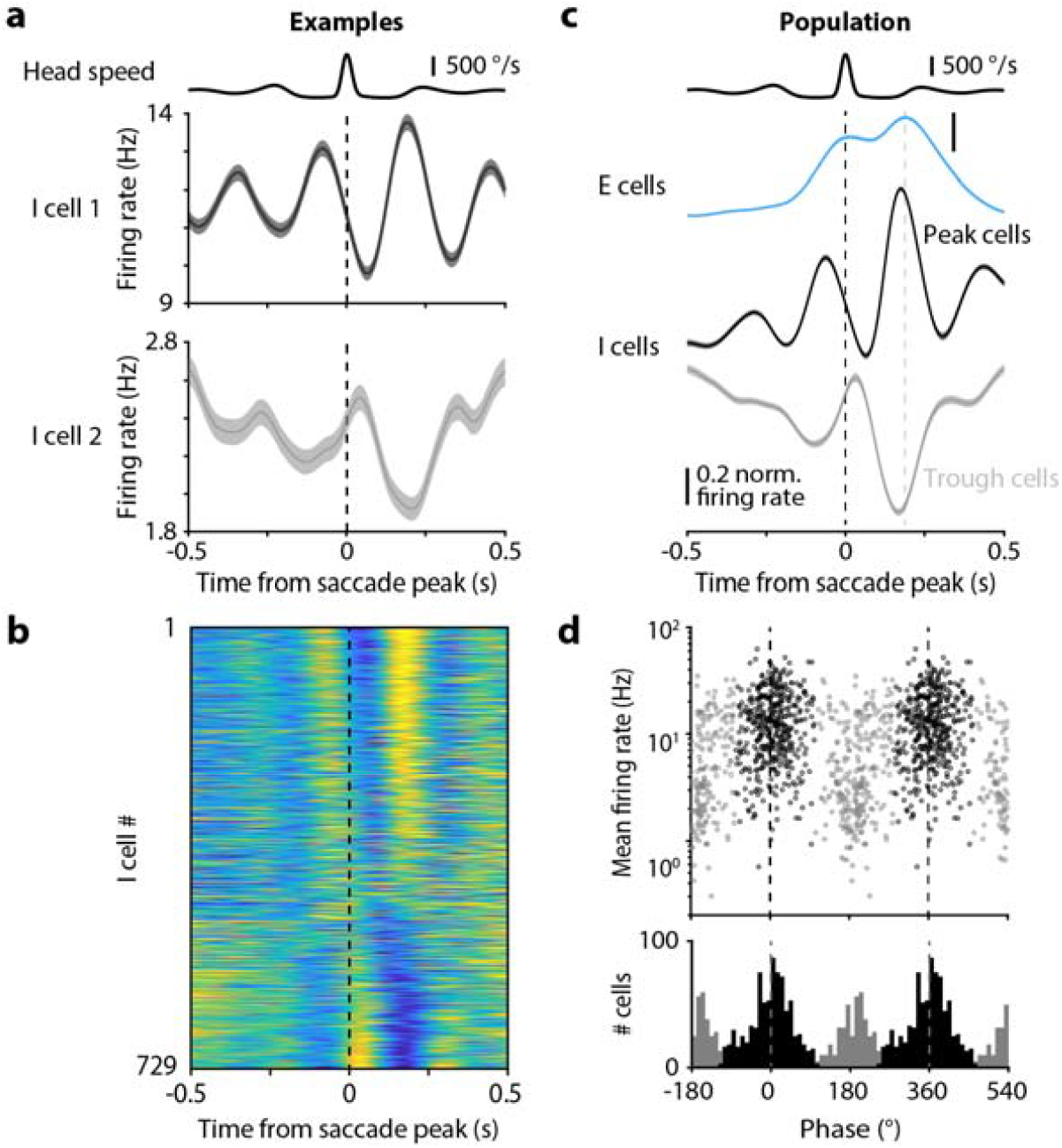
Inhibitory dynamics during saccades. **(a)** Activity of two inhibitory (I) cells, averaged across all saccades regardless of target, otherwise plotted as in Fig. 3a. **(b)** Responses of all I cells, sorted by the instantaneous Hilbert phase measured at 187 ms. Each cell’s response is normalized from minimum (blue) to maximum (yellow). **(c)** Responses of excitatory (E) cells and two types of I cells – Peak and Trough, classified according to the phase at 187 ms. Dotted lines indicate 0 and 187 ms – the time of the second peak in excitatory cell firing. **(d)** Distribution of phases and mean firing rates (across the entire session) for all I cells. Cells show a clear bimodality across the population.

Because saccades followed each other in quick succession, firing rates of I cells contained multiple peaks (Fig. 4a-c). For example, Peak cells produced the largest firing peak at 180 ms after the saccade. But since the median inter-saccadic interval was ∼270 ms (Fig. 1c), averaging across saccades produced multiple smaller versions of the same peak spaced roughly by that interval (e.g. at -90 and 450 ms). Owing to the multiple peaks, firing rates appeared to oscillate. Since saccades were irregularly timed, this was not a true periodic oscillation. Rather, the firing of I cells should be considered a quasiperiodic oscillation entrained by saccades. In this oscillation, Peak and Trough cells fire out-of-phase with each other, and are both phase-shifted relative to the E cells. Note that E cells did not show oscillatory firing because they had stronger spatial tuning than I cells. An E cell firing during one saccade was unlikely to respond to either the preceding or the following saccade because those saccades were toward different spatial targets. In summary, quasiperiodic saccade-related activity in chickadees has a major feature in common with theta oscillations in rodents: subtypes of interneurons that fire at different phases relative to each other and to the excitatory population.

## DISCUSSION

We have uncovered a critical role of vision in remotely driving place representations. Previously, gaze responses have been studied extensively in the primate hippocampus^9–14^. Considered without regard for place, the chickadee cells we recorded resemble monkey “view cells” – neurons with localized gaze responses that are invariant to the location of the animal. This similarity is notable because foveal vision in birds and primates evolved independently^4^ and depends on different movement strategies (head saccades vs. eye saccades). Localized gaze responses therefore appear to be fundamental to hippocampal function across these highly visual, but phylogenetically distant species. Yet prior to our work, there was a major missing link between these gaze responses and the well-studied spatial representations in the hippocampus.

The main reason for this gap is that experimental primates are usually stationary, and technical challenges discourage their recordings during movement. Only a few recent studies have managed to track gaze in navigating monkeys, either in virtual reality or with head-mounted devices^11–14^. These studies have not found the same close correspondence of place and gaze responses that we observed in chickadees. They have also reported only modest place and gaze selectivity, compared to the robust firing fields in chickadees. One issue is sampling: in these studies, monkeys rarely view and visit the same locations (most gazes are at distal landmarks rather that the floor). Another issue is that much of the monkey gazing behavior is passive, rather that deliberately directed at spatial goals. In contrast, our visual search task forced chickadees to gaze directly at their navigational targets, and ensured that they were attending to these targets. The motivational or attentional state could have a major effect on the hippocampal signals. Finally, our task design allowed cleanly separating periods of visual search from periods of navigation. Such analysis proved to be critical, since the hippocampus encoded gaze and place separately during these non-overlapping periods of time. Future experimental designs and analyses might uncover more similarities between birds and primates.

Remote activity of place cells in our data might also have a connection to findings in rodents. Rodent hippocampal activity often encodes places that are different from the actual location of the animal. This kind of remote coding famously happens during rest^43^, but can also occur during active behavior – especially when an animal is making navigational choices^30,31^. In these cases, hippocampal activity can represent potential future places and is hypothesized to influence navigational decisions. It is unknown what role vision plays in driving these remote activity patterns. Laboratory rodents are visual and move their eyes to control the field of view, but they lack a fovea and generally do not align their eyes precisely with visual targets^5,7,8^. Although eye position affects activity in their hippocampal circuit^44^, it is unknown whether this activity encodes gaze. A reasonable hypothesis is that rodents vicariously attend to distant visual targets, and that such attention drives remote coding in the hippocampus. In most behaviors, testing this hypothesis may not be possible by eye tracking alone. However, in some rodent behaviors like hunting, the precise visual target is known^8^. Hippocampal recordings in these behaviors will be informative for comparison with our results in chickadees.

Another intriguing connection of our work is to theta oscillations. Theta is important for several kinds of temporal coding in the hippocampus. In rodents, different molecular and morphological classes of interneurons fire at different phases of theta^40^. Their timing is essential for coordinating the firing of excitatory cells and for mechanisms of synaptic plasticity^41,42^. On each theta cycle, the hippocampus also switches between states dominated by external inputs and by internal connections^45^. This process might enable to hippocampus to fluctuate between different functions, such as memory storage during one phase of theta and memory recall during another phase. All these theories of theta are hard to reconcile with the fact that other species lack, or at least have greatly reduced theta^16,20,46^. We demonstrate a potential solution: temporal patterns of hippocampal activity can instead be paced by irregular saccades, forming a quasiperiodic oscillation. Animals might store and recall spatial memories (including food cache memories in chickadees) on different phases of the saccadic cycle. This fluctuation could be coordinated by specialized inhibitory cells – conceivably even cells homologous to those found in mammals.

Just like saccades synchronize hippocampal activity with incoming visual information, theta can synchronize activity with active sensory processes like whisking, sniffing, and stepping^47–49^. Saccadic modulation and theta might therefore be analogous processes, each adapted for the sensory behaviors of a particular animal species. Our results are consistent with work on the hippocampal LFP in bats, which is aperiodic^50^. They might also be consistent with primate data^16,51,52^ – though confusingly primates seem to retain some theta in addition to saccades. Monkey recordings so far have shown some differences in timing during saccades, including between E and I cells^14^.

Chickadee data allow us to formulate a general idea about hippocampal function. We suggest that hippocampal activity encodes the place that an animal is currently attending to. For a ground-foraging rodent, this place is almost always directly in front of the nose. For a highly visual animal like a bird or a primate, this place is usually at a distant visual target. In both cases, behavior forces some exceptions: rodents temporarily attend to distant targets to make navigational choices, while a moving bird may attend to the path directly ahead. As a result, both local and remote coding are present in the hippocampus, albeit in amounts that vary across species and behavioral tasks. The strength of our visual foraging task is that it required chickadees to use both modes of attention, and to switch between them at experimentally well-defined moments in time. Our results link disparate hippocampal phenomena like place coding, gaze coding, remote place activation, saccadic timing, and theta. They suggest how the hippocampus can simultaneously perform local functions (e.g. forming a new spatial memory when storing a nut) and remote functions (e.g. recalling that memory from a completely different place).

## ACKNOWLEDGEMENTS

We thank S. Hale for technical assistance; Aronov lab members for helpful discussion; the Black Rock Forest Consortium, J. Scribner and Hickory Hill Farm, and T. Green for field work support; and J. Kuhl for illustrations. Funding was provided by the NIH BRAIN Initiative Advanced Postdoctoral Career Transition Award (K99 EY034700, HLP), New York Stem Cell Foundation - Robertson Neuroscience Investigator Award (DA), Beckman Young Investigator Award (DA), NIH New Innovator Award (DP2 AG071918, DA), and the Howard Hughes Medical Institute (DA).

## Author contributions

H.L.P. designed and performed experiments, analyzed data, and wrote the paper; D.A. designed experiments, analyzed data, and wrote the paper.

