## Supplemental information for "Remote activation of place codes by gaze in a highly visual animal"

### METHODS

#### Subjects

All animal procedures were approved by the Columbia University Institutional Animal Care and Use Committee and carried out in accordance with the US National Institutes of Health guidelines. Subjects were 8 adult black-capped chickadees (*Poecile atricapillus*), 5 male and 3 female, collected from several sites in New York State using federal and state scientific collection permits. Of these, 7 were used in the Random task, 3 were used in the Blocked-trial task, and 3 were used in the All-to-all task, with some birds used in multiple tasks. Experiments were conducted blind to sex because chickadees do not have noticeable sexual dimorphism. Sex was determined post-hoc. During experiments, birds were singly housed on a “winter” light cycle, 9h:15h light:dark. Primary wing feathers were trimmed to prevent flight.

#### Head tracking and gaze estimation

To determine whether gaze could be estimated from head movements alone, and to estimate the direction of that gaze, we first conducted behavior-only calibration sessions for each bird. In these sessions the bird sat on a single perch and both head and eye movements were measured simultaneously (Fig. 1). For these sessions, we used the same behavioral arena as for the full task described below, but configured the floor with a single perch near a dish of seed fragments to encourage perching in one place.

We tracked head position using an infrared-based motion capture system (Qualisys Miqus cameras and Qualysis Track Manager software; Qualysis AB.) consisting of four specialized infrared cameras recording at 300 frames/s. The motion capture system tracked a 3D printed arrangement of 5 reflective markers (“head markers”, 3M Scotchlite 7610 Reflective Tape) affixed to an implant on the bird’s head using neodymium magnets and a 3D printed kinematic mount (Fig. 1a). Marker positions were calculated in a “**world**” reference frame anchored by known landmarks in the arena.

Eye position was tracked using a custom dual camera video-oculography system based on existing techniques that do not require cooperation of the subject for calibration^1,2^. The video-oculography system consisted of two cameras positioned 4 cm apart (Blackfly S BFS-U3-27S5M; lens Edmund Optics 59871 25mm/F1.4; visible light-blocking filter), and two infrared light sources positioned 11 cm apart (850 nm; Mouser 416-LST101H01IR0101). All cameras and light sources aimed at the bird sitting on the perch.

Before eye position could be recorded, there was a three step process to calibrate the combined head- and eye-tracking system. First, we determined the relative positions of the two cameras and their lens parameters using a checkerboard grid and the MATLAB computer vision toolbox (MATLAB, R2021a). This step defined a “**video-oculography**” reference frame centered on the optical center of the first camera. Second, we determined the positions of the infrared light sources relative to this video-oculography reference frame by imaging their reflections in a front surface mirror. The position and orientation of the mirror plane was determined using a checkerboard affixed to its mirrored surface. Finally, we aligned the world and video-oculography reference frames by determining the position of three reflective markers in both systems simultaneously.

For every frame in which the eye was visible in both video-oculography cameras, we used a published algorithm^3^ to determine the pixel coordinates of the pupil and the reflections of the two IR light sources on the corneal surface. Using our camera calibration described above, we then converted these 2D pixel coordinates to 3D positions in the video-oculography reference frame. Finally, we estimate the center of corneal curvature using the position of the two corneal reflections and the positions of the IR light sources relative to the eye and the cameras. We defined the position of the eye as the center of corneal curvature, and defined the optical axis of the eye as the vector pointing from the center of corneal curvature to the pupil center. MATLAB code for the eye tracking calibrations and analysis is available at <https://github.com/hpay/eyetrack-bird>). We applied several quality checks to discard frames in which the eye tracking failed.

We next defined a “**head**”-centered reference frame, which was applied to the calibration session as well as all experimental sessions. The position of the head (origin) was taken as the midpoint of the two eyes, averaged across frames. The x axis passed through the two eyes (right positive), the y axis passed through the midpoint of the two eyes and the tip of the beak (beak positive), and the z axis pointed up. We determined the orientation of the eye relative to the head in this reference frame. In Fig. 1d, we subtracted the mean horizontal and vertical angle of the eye from each data point to show the range of eye movements from rest. For the discrete visual search task described below, we used the mean position and vector of each eye in the head-centered reference frame to estimate the directions of gaze.

#### Behavioral experiments

All experiments were conducted in an enclosed square arena, with a central open space 61 cm on each side, surrounded by a 2.5 cm boundary interrupted by corner posts. The walls, floor, and ceiling were black, with ~15 cm diameter bright shapes (yellow circle, pink star, blue pentagon, and green tree) positioned on each wall, centered ~30 cm above the floor. The arena was illuminated from above. White noise was played in the background to mask inadvertent room noises.

Five feeder modules were positioned in the configurations described below for each task. Each module consisted of three concentric circles: a 3D printed perch (50 mm outer diameter, 30 mm inner diameter, 6.25 mm total height above arena floor), surrounding a raised ring of LEDs (eight DotStar LEDs per ring, Adafruit Industries) mounted on a custom PCB behind a diffuser (19 mm OD, 13m mm ID, 5.25 mm height above arena floor), surrounding a motorized feeder (11.6 mm opening diameter) that dispensed tiny sunflower seed fragments (~1.5 mg each) from a cup (4 mm deep). This arrangement ensured that the bird could not see into the feeder from a different perch, given a vertical head position of 54 ± 6 mm (mean ± standard deviation, n = 58 sessions).

In the Random and Blocked-trial tasks described below, the bird was encouraged to remain on the paths between the central and outer perches by a rubber surface restricted to an X shape, with each arm 7.5 cm wide. The rest of the arena was covered by a slippery ultra-high molecular weight polyethylene surface. For the All-to-all task, the rubber surface covered the entire arena, but birds still preferred to hop directly between perches.

To motivate food consumption, birds were deprived of food for 1-3 hours from waking (at the beginning of the light-on period of the day) until the start of the experiment. Birds were weighed daily before the experiment to ensure stable weight. Sessions typically lasted 1 hour. Birds typically underwent 3–6 habituation sessions, some conducted before surgery and some afterwards. Wired electrophysiological recordings began after these sessions. Weight from the implanted recording device and the cable was partially offset by a thin strand of fiber extracted from an elastic string (Linsoir Beads, Crystal String).

The light and feeder states were controlled in real time by the animal’s behavior. We tracked behavior at 300 frames per second using the reflective head markers and the calibrated motion capture system described above. The head marker coordinates were streamed from Qualysis Track Manager to MATLAB using the software interface QTM Connect for MATLAB (Qualysis AB). Then, the saved calibrations for each bird were used to determine head position and gaze vectors for each eye. In preliminary behavioral experiments, we found that birds typically directed their gaze towards targets along a vector slightly below the optical axis of the eye (Fig. 1h, *left*). We therefore rotated the estimated gaze vectors for each eye downwards by 5° during real-time tracking. The bird’s behavior controlled the experimental flow via Matlab code as described in detail below. Finally, Matlab sent serial commands to an Arduino Mega to change light and feeder states.

In all tasks, seed retrieval was detected when the bird’s beak tip was within 1 cm of the central site (if light was on but feeder was closed: 8 cm; pre-training of uncalibrated birds: when the bird’s head was within 5 cm of the central site). The feeder remained open for a fixed time, T_open_. We gradually reduced T_open_ from 20 s down to 1-2 s during pretraining, with the exact value chosen such that each bird had enough time to make only one beak poke. After T_open_, the feeder closed and the light turned off. Feeder opening and closing occurred smoothly over a total duration of 1 s.

In the **Random task**, there were five identical sites arranged in an X shape. Each outer site was located 34 cm from the central site. In this task, every session started with the central site (“Center”) illuminated (“turned on”) and its feeder open. Next, one of the four outer target sites was pseudorandomly selected as the “Target”. We ensured that the same target did not occur twice in a row, and that each target was chosen a roughly equal number of times per session. During some pre-training sessions (Fig. 1g), the Target turned on after a random delay (no more than 5 s). For the remaining experimental sessions, the Target only turned on when the bird was sitting at the Center and gazing towards the Target within a threshold of 10-20° of angular deviation for at least 3 time points. The on The median latency from the time of peak saccade velocity to the time of light onset was 40 ms (n = 8973 trials; Fig. 3e), and the median time to fixation onset was 33 ms. Both eyes could trigger the Target to turn on. If the bird visited any outer site before triggering the Target with gaze, the Center turned on and the program waited for the bird to return to the Center. The Target turned on every time the gaze trigger was activated. The feeder actually opened with a probability chosen manually based on the bird’s behavior (50 – 100%) in order to maintain motivation and increase the number of trials.

After the Target turned on, there were two variants of what happened next. In the “stable” variant of the task, the Target stayed on until the bird visited the feeder. In the “transient” variant, the Target only remained on while gaze was fixated at the target, but turned on again when the bird approached the site (within 8 cm). For the Random task, both task variants were used and pooled, since the analyses did not depend on the state of the light after initial detection. For the Blocked-trial task, all sessions were transient. For the All-to-all task, which was harder for birds to learn, all sessions were stable.

After the Target turned on, the program waited for the bird to visit the Target and, if the feeder actually opened, retrieve a seed. Incorrect site visits were indicated by turning the Target off and the Center on, and requiring the bird to visit the Center before the Target would turn on again. When the bird retrieved a seed from the target feeder, Target turned off and the Center turned on (with a low probability of reward, 5-25%). We considered a single trial to consist of one correct outward dash toward the Target and one inward dash toward the Center. Thus, this task elicited self-paced but structured center-out visual search behavior, with many dashes and saccades towards the same four outer sites.

The Blocked-trial task (Fig. 3f) had an identical physical arrangement. In contrast to the Random task, however, the Target was not chosen randomly, but instead was repeated six trials in a row. For the first trial in a block, the Target always turned on every time the bird’s gaze was detected. To increase the number of trials in each condition, only the contralateral eye could trigger gaze in this task. (For birds that were trained on both the Random and the Blocked-trial task, the Blocked-trial task was always run after the Random task to avoid introducing any bias in eye usage in the Random task.) For the second through sixth trials in a block, two of the trials were pseudorandomly chosen to be “Catch” trials. On a Catch trial, the first time the bird’s gaze was detected towards the Target, the Target stayed off. On all subsequent gaze detections, the Target turned on. The Catch trials were balanced such that over a session, there was a nearly equal number (±1) of Catch trials at each position within a block. In this task, the median latency from peak saccade velocity to fixation onset was 33 ms, and the median latency to light onset was 40 ms, as in the Random task.

The All-to-all task (Fig. S3) had five sites arranged in a pentagon. There was no Center site, and the Targets were chosen pseudorandomly in an all-to-all order. Sequences requiring the bird to visit three adjacent sites in a row were excluded from the pseudorandom assignment, and Target pair counts were balanced within a session. Incorrect visits were indicated by turning off the current Target and turning on the previous Target, requiring the bird to return to the previous site before activating a gaze trigger or feeder opening. Both eyes could trigger the Target to turn on.

#### Electrophysiological recording

We developed a light-weight system for chronic recording during free behavior (REF Selmaan). Neural activity was recorded using a 64-channel silicon neural probe (Cambridge NeuroTech, H5 or H6 ASSY-236). A three-part 3D printed housing system secured the headstage and protected the probe.

Signals were amplified, multiplexed, and digitized at 30,000 Hz using a custom PCB containing a wire-bonded RHD2164 chip (Intan Technologies, LLC). Intan RHX Data Acquisition Software (Intan Technologies, LLC) recorded the neural data simultaneously with the time of each video frame from the head tracking system, and the times of light or feeder changes from the behavioral control system. Digital signals were transmitted from the bird to a computer interface board via a 12-conductor SPI cable (Intan technologies, LLC, C3213), passed through a motorized commutator (Doric Lenses, Inc., AERJ_24_HDMI).

To minimize degradation of neural signals over time, the probe contacts were left embedded in a silicone gel covering the brain when not in use. The probe was inserted to the desired depth 30 minutes before recording and retracted at the end of each session using an aluminum nano-drive (Cambridge NeuroTech).

The entire assembly was 1.1 g (0.1 g probe, 0.40 g headstage and connectors, 0.28 g drive, 0.32 g housing); cement added ~0.2 g more and the head tracking markers added 0.14 g more.

#### Surgery

Surgery was conducted using a two-step procedure, largely as described in (Chettih et al 2024). In the first step, a dummy implant with a removable cap was affixed to the skull. The bird was allowed to recover and removable weights were gradually added. In the second step, the craniotomy was made and the probe was implanted.

During the first step, birds were anesthetized using 1.5% isoflurane in oxygen. Feathers were removed from the surgical site, and the bird was placed in a modified stereotaxic apparatus using ear bars and a beak clamp. The head was tilted such that the angle of the groove at the base of the upper mandible of the beak was 65° relative to the horizontal, corresponding to an angle of 30° between the bite bar and the horizontal. A silver ground wire (0.005” diameter) was implanted in the contralateral hemisphere, posterior and lateral to the hippocampus and 1 mm below the surface. The location of the probe craniotomy was marked on the surface of the skull: 3.02 – 4.05 mm anterior to lambda, 0.5 – 0.73 mm lateral to the midline. Most microdrives were implanted into the left hippocampus, with two microdrives implanted in the right hippocampus. The tilt of the head was adjusted to that the 3D printed base would sit flat on the skull when centered over the craniotomy. A short base unit was cemented over the planned craniotomy (RelyX Unicem, 3M). A removable 3D printed cap was attached to the base unit. After the surgery, buprenorphine (0.05 mg/kg) was injected intraperitoneally and the bird was allowed to recover for 1.5-2 weeks while weight was monitored. After at least 5 days, a 1 g weight was added to the dummy cap.

During the second step, the bird was anesthetized as above and injected intraperitoneally with dexamethasone (2 mg/kg). The cap of the dummy implant was removed and the remaining base unit and skull were cleaned with 70% ethanol. A craniotomy and durotomy were performed covering a 1x1 mm area centered on the coordinates given above. A 3D printed biocompatible resin insert with a small central slit (0.4 x 0.1 mm) was inserted into the craniotomy site so that it pressed gently on the brain surface and was cemented to the skull. A silicon probe mounted on a drive was then moved into place, tilted laterally by 10°-15° to target the medial hippocampus. After insertion of the probe was checked, the space above the craniotomy was filled with a protective layer of silicone elastomer (Dow Corning 3-4680 Silicone Gel). The probe was advanced to sit in the elastomer and the drive was cemented into place. The protective outer housing and the headstage were secured over the probe. The top surface of the outer housing contained a hole to access the drive screw, and a kinematic composed of two small neodymium magnets and 3D printed features to allow reliable re-positioning of the reflective markers during behavioral sessions.

#### Histology

After completion of all experiments for each bird, the silicon probe was left in place overnight. Birds were given an overdose of ketamine and xylazine and were then perfused transcardially with saline followed by 4% formaldehyde. Brains were extracted and stored in 4% formaldehyde, then cut into 100 μm-thick coronal sections. Brain sections were stained with fluorescent DAPI. The position of each electrode relative to the boundary of the hippocampus was estimated by measuring the distance from the surface of the brain to the lateral ventricle along the electrode track. This measurement was used to exclude recorded cells that were likely outside the hippocampus.

#### Spike sorting

All analyses were conducted in MATLAB unless noted. Spike sorting was conducted using Kilosort2.0^4^. Default settings were used, except that the high-pass filtering cutoff was set to 300 Hz and there was no minimum firing rate for good channels. A total of 25 sessions were manually curated in Phy (Python), including 15 in the current dataset and 10 from previous pilot experiments. During manual curation, the automatic labels were edited as needed to mark units as “good”, “mua” (multiunit activity), and “noise”. The remaining 55 sessions were automatically curated by applying several criteria. First, we calculated the spatial extent of each unit along the probe, as well as the cluster contamination rate determined by Kilosort, and excluded units that passed a threshold for each.

Second, we identified putative excitatory and inhibitory neurons by applying a Gaussian mixture model (GMM) to the following four electrophysiological characteristics (Fig. S5): spike rate (log transformed), spike width, spike asymmetry, and derivative peak-trough ratio. Spike width was calculated as the time from the trough of the average spike waveform to the subsequent peak. Spike asymmetry was calculated as the relative height of the two positive peaks flanking the trough. Derivative peak-trough ratio was the log-transformed ratio of the peak amplitude to the trough amplitude of the waveform derivative. We fit the GMM on the manually curated sessions to classify cells into two clusters corresponding to putative excitatory and inhibitory neurons. We then applied the GMM to all sessions and excluded units that exceeded a distance threshold from either of the two clusters. A small number of neurons that were intermediate between the two clusters were labelled as unclassified neurons and were not considered further. Cells with fewer than 500 spikes were excluded (457/3115 cells). Custom code to run spike sorting and process results is available at: <https://github.com/hpay/spikesort-hp>.

Inhibitory neurons were further clustered into two groups on the basis of their average saccade-aligned activity (see Main Text). Mean firing rate was 1.3 Hz, 14.0 Hz, 7.1 Hz in the putative excitatory, Late inhibitory, and Early inhibitory clusters, respectively; spike width was 0.51 ms, 0.27 ms, and 0.33 ms; and peak amplitude asymmetry was -0.04, 0.55, and 0.56.

#### Behavioral analysis

The position and orientation of the head were tracked using the motion capture system as described above. Linear head speed was calculated by differentiating the x, y, and z position of the head and calculating absolute speed as $\sqrt{\Delta x^{2}+\Delta y^{2}+\Delta z^{2}}/\Delta t$. Angular head speed was calculated by measuring the angular difference in 3D orientation of the head between adjacent frames and dividing by $\Delta t$. Both linear and angular speed were then low-pass filtered with a Butterworth filter with a cutoff frequency of 25 Hz.

##### Hidden Markov Model

To label behavioral states, we implemented a Hidden Markov Model (HMM) based on previous work^5^. The HMM predicted whether the bird was saccading, fixating, or feeding at each of the five sites, or whether it was dashing between two pairs of sites. The HMM had a Gaussian observation model, representing the likelihood of observing the position, orientation, and speed of the bird’s head at each time point given each state. The most likely state was computed using the Viterbi algorithm. Input data consisted of linear speed, angular speed, and head position and tilt. Position was transformed to represent proximity to sites or lines between sites. Mean parameters of the Gaussian model were fit iteratively for each session, while the variance parameters were fixed. The transition probability matrix was fixed.

We refined the output of the HMM as follows. This refinement step was prompted by our observation of common misclassifications of dashes, saccades, and fixations. First, we cleaned up misclassified states. Any “saccades” immediately preceding feeding were combined with the feeding state. Any “saccades” immediately following a dash were combined with the dash state. Any “feeding” after a dash was combined with the dash. Second, we refined the endpoints of saccades and dashes (see function in GitHub repository refineHMM.m). The start and stop times of each saccade were refined by applying velocity (400°/s) and acceleration (5000°/s^2^) thresholds. The stop time of each dash was also refined by applying velocity (150 mm/s) and acceleration (3000 mm/s^2^) thresholds.

Sessions with fewer than 5 dashes or 5 saccades with each eye to each outer target were excluded from further analysis. Unless otherwise noted, all analyses were conducted on saccades that were immediately followed by a fixation. We used the time of peak saccade velocity as the alignment point for all analyses (the “time of the saccade”).

##### Analysis of gaze strategies

To calculate the time course of gaze strategies shown in Fig. 2g and Fig. S1h, a histogram of head orientations relative to the target was first calculated for each bird, and the peak density near the beak and near each eye (averaged across eyes) was taken as the ideal vector for frontal and lateral gaze, respectively. For each saccade in the sequence, we then determined the angular distance between either frontal gaze or lateral gaze with either eye and the target of the upcoming dash. Angular distances less than 20° counted as a “hit”, and the probability of a hit for either frontal or lateral gaze across all saccades was plotted.

#### Neural analysis

##### Place and gaze responses

To construct place map examples (Fig. 2c), we included position data during dashes plus a ±1 s window before and after each dash. Spatial information was calculated using a range of delays between spikes and behavior, and the best offset was chosen. The arena floor was divided into 40×40 bins in which spike counts and occupancy time were calculated. The resulting matrices were each smoothed with a 11×11-point Hamming window then divided to yield mean firing rate.

The gaze maps (Fig. 2d) included data during gaze fixations made while at the Center site. Gaze from the specified eye was projected along the optical axis measured during the calibration session from that bird. A cone with a radius of 10° was projected on the floor. Spike counts and occupancy times were calculated, smoothed with a 9×9-point Hamming window, and divided to yield mean firing rate in each bin.

We quantified the degree of spatial tuning for each neuron for both place and gaze. We quantified place tuning by calculating the information about site identity conveyed by the cell’s firing during dashes towards each of the four outer sites. We calculated information according to:

$$I= \sum_{x} \frac{\lambda\left( x \right)}{\lambda}\log_{2} \frac{\lambda\left( x \right)}{\lambda}p(x)$$

where *I* is the information rate in bits/spike, 𝑥∈{1,2,3,4} is the site identity, *p*(𝑥) is the probability that the bird visited site 𝑥, *λ*(𝑥) is the mean firing rate in a ±1 s window centered on the end of each dash to site 𝑥, and $\lambda=\Sigma\lambda\left( x \right)p\left( x \right)$ is the overall mean firing rate across all included time periods^6^. A null distribution for each cell was calculated by shuffling the identity of the dash targets across trials and re-calculating spatial information for 200 repetitions. A neuron was considered a significant “place cell” if actual spatial information exceeded 99% of samples in the shuffled distribution. Spatial information was normalized for each cell by dividing actual spatial information by the mean of the shuffled distribution (Fig. 1g).

We applied the same procedure to quantify the degree of gaze tuning, but for spike counts within a window from -0.1 s to +0.3 s from the time of peak saccade velocity. Only saccades that landed on a target (within 20° of visual angle), and that occurred more than 0.5 s before the start of a dash, were included.

Firing rates were calculated over time by binning spikes at the frequency of the behavioral data acquisition (3.33 ms bins) and applying either a 100-ms or a 30-ms sigma Gaussian filter for dash or gaze responses, respectively.

Dash responses were summarized across the population (Fig. 2e) by calculating the average firing rate over time aligned to dashes to each of the four outer sites for each neuron. The peak of the response in a ±1 s window centered on the dash end was measured for each of the four sites. Saccade responses were summarized across the population (Fig. 2f) using a different approach, because the configuration of sites (separated by 90°) resulted in behavioral correlations between gaze with each eye to adjacent sites (angle between eyes 106 ± 5°, mean ± standard deviation, n = 8 birds). We estimated the separate contributions of ipsiversive and contraversive gaze to neural firing using a Poisson generalized linear model (GLM). Lasso regression was applied using the Matlab function lassoglm with alpha = 1 and lambda = 0.005 to discourage overfitting. The model was given by:

$$\log\left( E(\boldsymbol{y}|\boldsymbol{X}) \right)=\boldsymbol{\beta}^{\boldsymbol{'}}\boldsymbol{X}$$

where each element of $\boldsymbol{y}$ was a scalar $y_{i}$ containing the observed spike count within a window from -0.1 s before to +0.3 s after the time of peak saccade velocity for trial *i*. Each column of $\boldsymbol{X}$ contained the predictors $\boldsymbol{x}_{i},$ which were given by:

$$\boldsymbol{x}_{i}=e^{-\frac{\boldsymbol{\alpha}^{2}}{\tau^{2}}}$$

where $\boldsymbol{\alpha}$ was a vector representing the angular distance of each site from each eye’s gaze vector, contralateral (“C”) or ipsilateral (“I”) to the site of recording:

$$\boldsymbol{\alpha=}\left[ \alpha_{C}^{1}\alpha_{C}^{2} \alpha_{C}^{3} \alpha_{C}^{4} \alpha_{I}^{1} \alpha_{I}^{2} \alpha_{I}^{3} \alpha_{I}^{4} \right]$$

and $\tau$ was a length constant equal to 45°, determined by fitting the decay of neural activity across the population as a Gaussian function of distance.

For the All-to-all task, birds started each trial in a different location, so place codes for a preferred site could contaminate responses during gaze from that site to other non-preferred sites. To account for this, we subtracted baseline activity from both the dash and saccade responses in this task as follows. The neural response for dashes was calculated as the mean response in a ±0.5 s window centered on the dash end, minus the mean response from -2 s to -1 s. The neural response for saccades was calculated as the mean response in a 0 s to +0.3 s window aligned to the saccade, minus the mean response from -1 s to -0.2 s. Responses below baseline were included in the analysis, but the color plots are cropped at zero. All included cells had at least four saccades and four dashes for every source-target pair (20 total permutations).

We determined the selectivity of each cell for place or gaze at a single site by comparing responses for the preferred site to the next-most-preferred site. Responses for dash and gaze (either peak rates for dash, or model coefficients for gaze) were first normalized by dividing by the response for the preferred site. A selectivity index was then calculated:

$$I_{select}=r_{1}-r_{2}$$

where $r_{1}$was the normalized response for the site with the largest response and $r_{2}$ was the response for the site with the second largest response.

In Fig 3c-d, cells are sorted by their difference in firing during the early and the late responses relative to head saccades. To define early and late responses, we calculated the mean firing aligned to saccades for each cell. We then averaged these responses across all excitatory cells with strong selectivity (>0.5) of both place and gaze responses for the same target, and finding the two peaks in the average response (Fig 4c, *blue*). For each cell, we then found the peak firing rate within a ±50 ms window centered around each of the two peaks. Cells were sorted by the difference in peak rate during these two windows.

##### Analysis of interneuron firing

We categorized the pattern of interneuron firing by first calculating the mean firing rate for each cell across all saccades. Interneurons had some selectivity, but were generally active for all saccade targets. We then performed a Hilbert transform on the mean firing rate, and stored the instantaneous phase of the response at 187 ms after the saccade peak (the time of peak firing in excitatory cells, Fig. 4c). We performed circular k-means clustering on the instantaneous phases to classify interneurons into two groups.

#### Statistical analysis

All confidence intervals given in the text and figures are mean ± standard error of the mean unless otherwise specified.

### SUPPLEMENTARY FIGURES

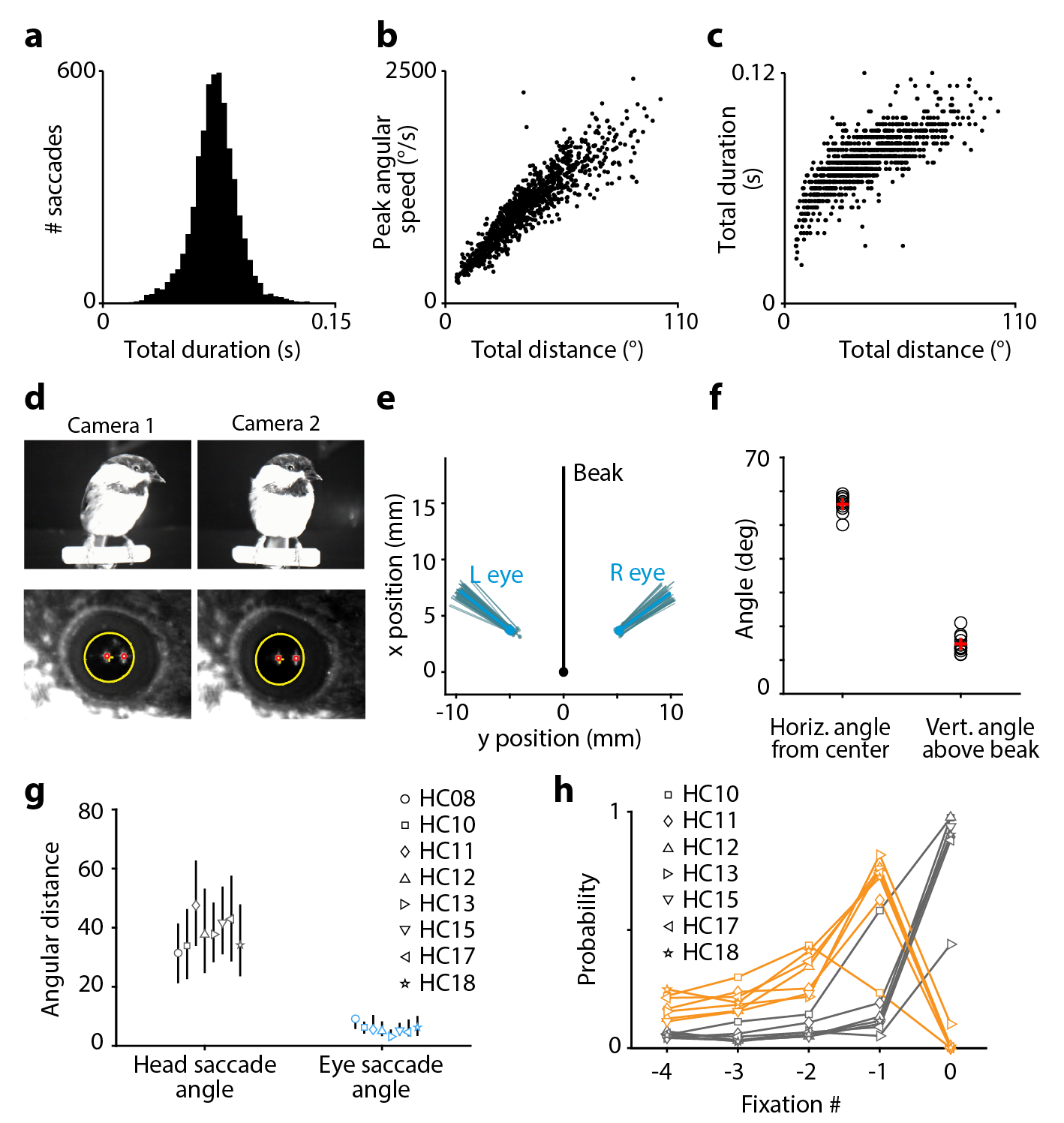

**Extended Data Figure 1. Properties of head and eye movements in chickadees
(a)** Head saccade durations for an example session. **(b)** Relationship between angular distance travelled by the head and the peak angular speed during head saccades for the same session. **(c)** Relationship between distance travelled and saccade duration. Both (b) and (c) show strong correlations, illustrating similarities to the eye saccade main sequence in primates^7^. **(d)** Video-oculography using two infrared light sources. The chickadee is perched close to two cameras, which acquire video frames from slightly different angles (*top*). The pupil (yellow) and the corneal reflections of the two light sources (*red*) are detected in the videos. **(e)** Position of the eye (*dots*) and the orientation of the optical axis (*lines*) relative to the head. Projections onto the horizontal axis are shown. Gray: individual video frames for a single calibration experiment. Blue: average across frames. **(f)** Orientation of the optical axis across all birds (*black symbol*) and the average across birds (*red symbol*). **(g)** Angular displacement of the head and the eye during head saccades. Symbols indicate medians for each bird; lines indicate 25th and 75th percentiles. **(h)** Time courses of the two gaze strategies (*orange*: lateral; *black*: frontal). Data are shown as in Fig. 1g, but for all birds, averaged across sessions.

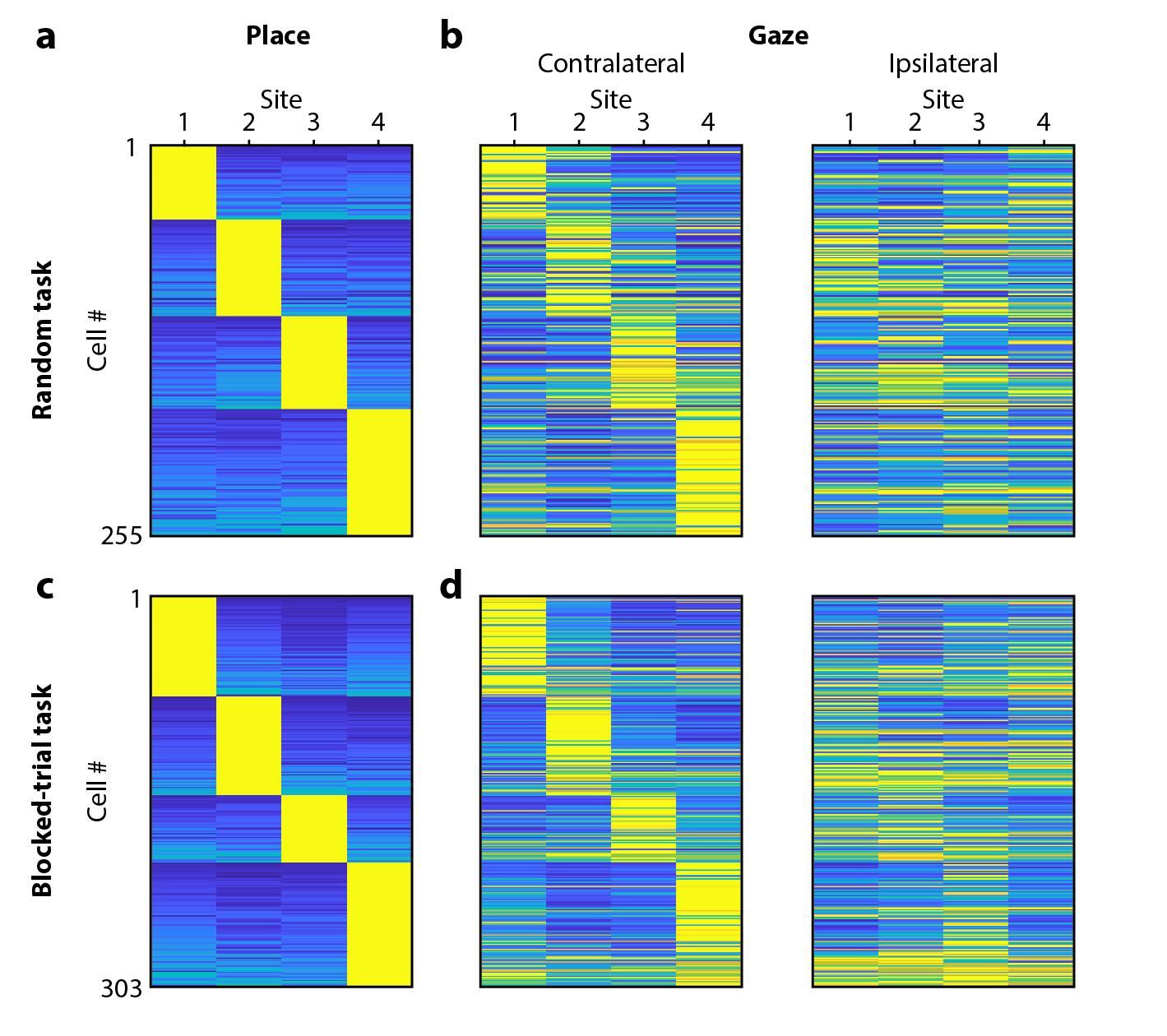

**Extended Data Figure 2. Comparison of place and gaze coding across tasks
(a-b)** Place and gaze coding, shown as in Fig. 2e-f, but only for cells recorded in the Random task, where the location of the rewarded target was chosen randomly on each trial. **(c-d)** Same, but only for cells recorded in the Blocked-trial task, where the same rewarded target was repeated for six trials in a row.

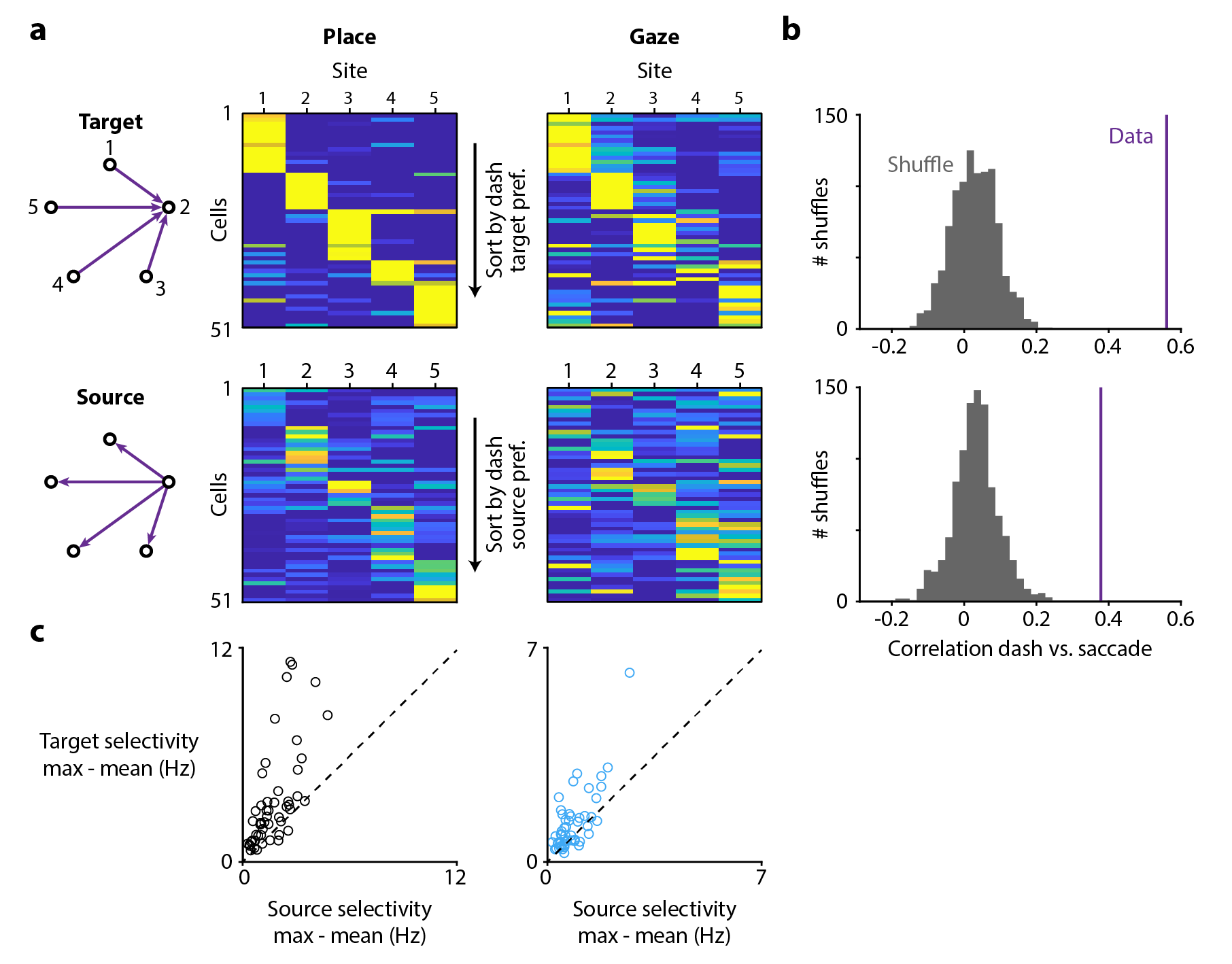

**Extended Data Figure 3. All-to-all task to disambiguate gaze target from the bird’s location (a)** Activity in the All-to-all task, where the chickadee was located at one of five sites and gazing at one of the other sites. After this visual search period, the chickadee dashed directly from one site to another. The location of the bird prior to the dash was the “source”, while the target of gaze and the endpoint of the dash was the “target”. Activity of each cell is shown during dashes (“Place”, *left*) and during gaze fixations (“Gaze”, *right*). Firing rates are calculated as a function of either the target site (*top*) or the source site (*bottom*). Included are cells with place and gaze selectivity (>0.67), either for the target or the source, and with baseline-subtracted response > 0.5 Hz. Each row is normalized from 0 (*blue*) to maximum (*yellow*) across target and source measurements, separately for place and gaze. **(b)** Correlation of the tuning curves for place and gaze (i.e. the rows of the matrices in (a)). *Top*: correlation of target tuning curves; *bottom*: correlation of source tuning curves. In both cases, correlations for actual data are higher than for a shuffled distribution, in which cell identities were scrambled. **(c)** Comparison of selectivity for target and for source. Selectivity was measured by subtracting the mean of the tuning curve from the maximum. For both place (*left*) and gaze (*right*), selectivity is higher for the target than for the source.

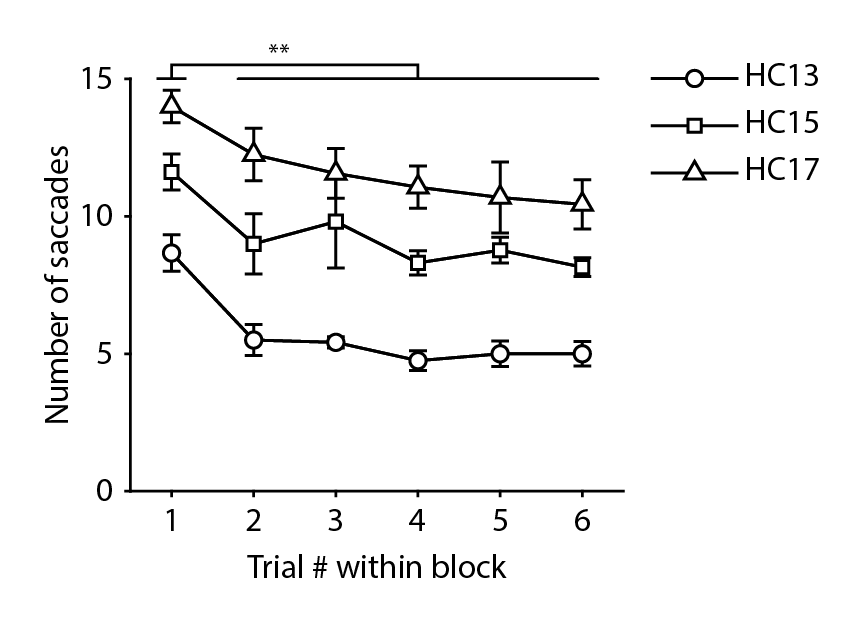

**Extended Data Figure 4. Chickadees understand the structure of the Blocked-trial task**
Number of saccades required by birds to find the correct target for different trials within each block. Chickadees take longer to find the target on the first trial (when they have no information about which target is rewarded) than on subsequent trials (all p < 0.01, two-sample t-test conducted separately for each bird). Note that when shifting gaze from one target to another, birds often make several intermediate saccades at other points in the environment.

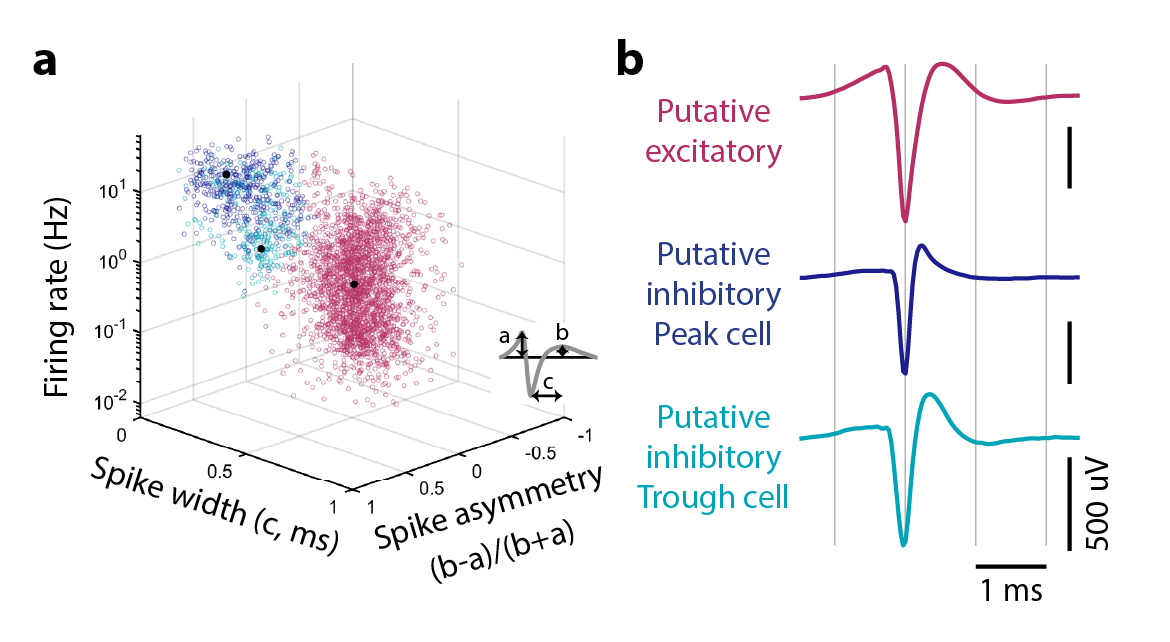

**Extended Data Figure 5. Classification of cell types in the chickadee hippocampus**
**(a)** All recorded units plotted according to their mean firing rate across the session and two features of spike waveforms shown in the diagram. These measurements separate putative excitatory (*pink*) from putative inhibitory (*blue* and *teal*) cells. Inhibitory cells are further classified by their responses during saccades (Fig. 4) into Peak (*blue*) and Trough (*teal*) types; these types show some systematic differences in firing rate and spike waveforms. **(b)** Average spike waveforms of three example cells, one from each of the categories shown in (a).
